## Supplemental material for "Imaging analysis of six human histone H1 variants reveals universal enrichment of H1.2, H1.3, and H1.5 at the nuclear periphery and nucleolar H1X presence"

#### SUPPLEMENTARY MATERIAL

This file contains the Supplementary Figure Legends and Supplementary Tables corresponding to the manuscript: ***Imaging analysis of six human histone H1 variants reveals universal enrichment of H1.2, H1.3, and H1.5 at the nuclear periphery and nucleolar H1X presence.***

#### SUPPLEMENTARY TABLES

**Supplementary Table 1. Oligonucleotides for semiquantitative PCR.**

#### SUPPLEMENTARY FIGURE LEGENDS

**Figure 1-figure supplement 1. Endogenous H1 variants immunofluorescence controls and colocalization with heterochromatin protein HP1. A)** To discard that the peripheral nuclear enrichment seen by immunofluorescence for some of the H1 variants is due to saturating antibody concentrations, we performed H1.3 immunofluorescence using serial dilutions of primary antibody. Graphs show immunofluorescence quantification signal of H1.3 using different antibody concentrations, as indicated. As illustrated in **Figure 1B**, four sections of an equivalent area and convergent to the nuclear center are created per each cell. Sections are named A1 to A4, from the more peripheral section to the more central one. H1.3 immunofluorescence intensity is measured in each area and expressed as percentage. n=30 cells/condition were quantified, and data was represented in violin plots. Statistical differences between consecutive sections are supported by paired t-test (***) p-value<0,001. Of note, a clear peripheral distribution was observed for H1.3 despite the antibody concentration used. **B)** Immunofluorescence of T47D cells stably expressing HA-tagged H1.0 was done using H1.0 and HA antibodies to evaluate co-localization between both antibodies signals. Panels show intensity profiles of H1.0 and HA antibodies along the arrows depicted. Scale bar: 5µm. A violin plot showing the Pearson correlation between H1.0 and HA signals in n=56 cells is included. The high overlap and correlation obtained between HA and H1.0 prove the ability of the antibody to recognize epitopes in different chromatin environments despite H1 location. They also prove that the nuclear peripheral enrichment observed for some variants is not due to the inaccessibility of the antibody to more central or occluded regions. **C)** Confocal immunofluorescence of HP1alpha (red), H1 variants (green), DNA staining (blue). Bottom panels show the intensity profiles along the white lines depicted in the merge panel. Correlation between HP1a and the corresponding H1 variant along the depicted line is indicated above the graphs. In the bottom profiles, gray arrows point to HP1a foci. Scale bar: 5µm. In **D) T**hree additional profiles per H1 variant are shown.

**Figure 2-figure supplement 1. H1 variants co-localization at confocal resolution. A)** Confocal immunofluorescence of the indicated H1 variants (green) with H1.0 (red) and

DNA staining (blue). Zoom-in insets highlight H1.0-peripheral enrichment territories. Scale bar: 5µm. **B)** Violin Plots showing the Pearson correlation coefficient (r) distribution of H1 variants with H1.0 in n=40 cells/condition. **C)** Statistical comparison of results shown in (B). ANOVA multiple comparison test revealed that significant differences exist between groups. Tukey multiple comparison test was used to compare the H1s-H1.0 r values distribution between different H1s. p-adjusted values are shown (***) p-adj< 0.001; (**) p-adj<0.01; (ns/non-significant) p-adj >0.05).

**Figure 2-figure supplement 2. H1 variants distribution patterns along mitosis phases.** Immunofluorescence of H1 variants and LaminA along the distinct mitotic phases. In all cases, DNA staining is also shown. **A)** Replication-independent H1.0 and H1X variants. **B)** H1.5 and Lamin A. **C)** H1.3 and Lamin A. **D)** H1.2 and Lamin A. Scale bar: 5µm

**Figure 2-figure supplement 3. H1 variants phosphorylation during mitosis. A)** Immunofluorescence of H1.2-pT165 and H1.4pT146. A Z-stack maximum projection is shown. Mitotic cells are marked by a red arrow. Scale bar: 10µm. **B-C)** Immunofluorescence of phosphorylated H1.2-pT165 (B) and H1.4-pT146 (C) along the distinct mitotic phases. DNA staining is also shown. Scale bar: 5µm

**Figure 3-figure supplement 1. H1 variants within LADs and nucleoli under basal and upon rDNA transcription inhibition. A)** Representative confocal immunofluorescence images of H1 variants (green), H3K9me2 (red) and DNA (blue). **B)** Immunofluorescence of H1X and DNA in T47D H1Xsh -/+Dox. A Z-stack of five consecutive Z planes is shown. Scale bar: 10µm. H1X immunofluorescence signal quantification (n=41 cells/condition) is shown and supported by paired-t-test. (***) p-value < 0.001. **C)** Immunofluorescence of NPM1, H1.4-pT146 and DNA in T47D cells. A unique central confocal Z plane (1Z) or the Z maximum projection (Zproj) are shown. Scale bar: 5µm. **D)** Immunofluorescence of H1X, H1.2p-T165 or H1.4-pT146 (green) co-immunostained with Nucleophosmin (magenta) and DNA (blue) under Untreated or Act-D-treated conditions. Scale bar: 5µm

**Figure 4-figure supplement 1. Super-resolution imaging of DNA upon different H1 KD conditions. A)** Representative images of DNA staining visualized with wide-field or super-resolution (SRRF) microscopy in a T47D nucleus. Bottom panels show a zoom-in of the perinucleolar region. Scale bar: 5µm (full nucleus) or 500nm (Zoom-in insets). **B)** Additional representative SRRF images of DNA staining in the different H1 KD conditions (multiH1, H1.2, H1.4 and H1X Dox-inducible shRNAs). In the bottom pannels, a zoom-in inset is shown to appreciate DNA pattern in both Untreated and Dox conditions. Scale bar: 5µm (full nucleus) and 500nm (zoom-in). **C)** Quantification of % DNA-free areas within n=20 cells per condition.

**Figure 5-figure supplement 1. H1 protein and mRNA complement across cell lines. A)** Immunoblot analysis of H1 variants in histones extracts from different tumoral and non-tumoral cell lines. Histone H3 is added as a nuclear control and Coomassie staining is shown. **B)** Coomassie staining quantification of histone extracts from cell lines shown in

(A). Graph shows the contribution of the three H1 Coomassie bands to total H1 content. Data is normalized by H4 band and represented as percentage. **C)** Immunoblot of H1 variants in histones extracts (2 or 10 µg) from six melanoma cell lines. Histone H3 and H4 are added as nuclear controls and Coomassie staining is shown. BRAF mutation status is indicated. **D)** Top panels show the ImageJ profiling of histones Coomassie shown in (C). The bottom panel represents the contribution of the three H1 Coomassie bands to total H1 content. Data is normalized by H4 band and represented as percentage. In cell lines lacking H1.3 and H1.5, the upper H1 Coomassie band corresponds to H1.4. **E)** Gene expression levels of H1 variants in different cancer cell lines were analyzed by RT-qPCR. Data is corrected by GAPDH and normalized by the corresponding genomic DNA amplification. Corrected expression data from all H1 variants is summed to calculate total H1 expression and represent values as percentage. Data is represented as pie charts or numerically collected in a table.

**Figure 5-figure supplement 2. H1 variants expression regulation by DNA methylation. A)** Scatterplots between H1 variants expression (Y-axis) and H1 gene methylation (X-axis) from NCI-60 public data. Gene expression from total RNA-Seq data is expressed in log2 (FPKM+1) while gene methylation is expressed as b value (β =0 is totally unmethylated and β=1 is totally methylated). Each dot represents a cell line from the NCI-60 panel. Pearson correlation coefficients (R) are shown. **B)** Boxplots show the DNA methylation of H1 genes in different cancer datasets from TCGA project. Gene methylation is expressed as β value (β =0 is totally unmethylated and β =1 is totally methylated). **C)** H1 variants expression levels in different cell lines under Untreated and aza-treated conditions were analyzed by RT-qPCR. Barplot shows relative expression of H1 variants upon aza treatment compared to Untreated condition, corrected by GAPDH and expressed as log2. SK-MEL-147 is added as a control cell line which expresses all H1 variants (except H1.1).

**Figure 5-figure supplement 3. H1 variants nuclear distribution in non-tumoral cell lines.** Confocal immunofluorescence of H1 variants (green) and DNA staining (blue) in 293T **(A)** or IMR-90 **(B)** cells. Bottom panels show the intensity profiles of H1 variants and DNA along the arrows depicted in the corresponding upper panel. Scale bar: 5µm. Of note, 293T do not express H1.5 and have very low levels of H1.0 **(see Figure 5-figure supplement 1A)**. Indeed, most 293T cells are H1.0-negative in immunofluorescence. Regarding IMR-90 cells (B), it is important to note that these cells have a characteristic DNA staining pattern with DNA foci. H1.2, H1.3, H1.5 and H1.0 variants are highly enriched in these DNA foci, including those in the periphery. Thus, these profiles support the enrichment of these variants at heterochromatin or low-GC regions, present, but not limited to the nuclear periphery. On the contrary, H1.4 and H1X are distributed throughout the whole nucleus, with H1X being highly present at nucleoli, but they are not enriched in these heterochromatic foci. This observation supports the two H1 groups, in terms of distribution, observed in T47D.

**Figure 5-figure supplement 4. Nuclear radial distribution of H1.4 and H1.0 across cell lines.** Graphs show immunofluorescence quantification signal of **A)** H1.4 and **B)** H1.0 in different cell lines. As illustrated in **Figure 1B**, four sections of an equivalent area and convergent to the nuclear center are created per each cell. Sections are named A1 to A4, from the more peripheral section to the more central one. H1 variants immunofluorescence intensity is measured in each area and expressed as percentage. Cell lines with a compromised H1 somatic repertoire are indicated. n=30 cells/cell line were quantified, and data was represented in violin plots. Statistical differences between A1-A2, A2-A3 and A3-A4 are supported by paired t-test (***) p-value<0,001; (*) p-value < 0.05; (ns/non-significant) p-value>0.05.

**Figure 5-figure supplement 5. Cell lines lacking H1.3 and H1.5 show high basal expression of repetitive elements in comparison with cell lines with a complete H1 somatic repertoire. A)** RT-qPCR of interferon stimulated genes (ISGs) and repetitive elements in different melanoma cell lines. Expression data is corrected by GAPDH and normalized by SK-MEL-147 cell line basal expression, which presents a complete H1 repertoire. SK-MEL-173 and IGR-39 cell lines show concomitant absence of H1.3 and H1.5 **(see Figure 5-figure supplement 1C**). **B)** RT-qPCR of ISGs in breast cancer cell lines. MDA-MB-231 cell line does not express H1.3 and H1.5**.** Expression data is corrected by GAPDH and normalized by basal expression in T47D multiH1sh Untreated cells. T47D multiH1sh+Dox cells, which are reported to trigger a high expression of multiple ISGs (ref Izquierdo-Bouldstridge et al, 2017 in the main manuscript) are also shown for comparison.

| | SENSE | SEQUENCE (from 5' to 3') |
|---|---|---|
| H1.0 | F | CCTGCGGCCAAGCCCAAGCG |
| | R | AACTTGATCTGCGAGTCAGC |
| H1.1 | F | CTCCTCTAAGGAGCGTGGTG |
| | R | GAGGACGCCTTCTTGTTGAG |
| H1.2 | F | GGCTGGGGGTACGCCT |
| | R | TTAGGTTTGGTTCCGCCC |
| H1.3 | F | CTGCTCCACTTGCTCCTACC |
| | R | GCAAGCGCTTTCTTAAGC |
| H1.4 | F | GTCGGGTTCCTTCAAACTCA |
| | R | CTTCTTCGCCTTCTTTGGG |
| H1.5 | F | CATTAAGCTGGGCCTCAAGA |
| | R | TCACTGCCTTTTTCGCCCC |
| H1X | F | CCCAACGATGTAGCGTTTTT |
| | R | AAGGCCGAGAGCCAATAGA |
| IFi27 | F | TGCTCTCACCTCATCAGCAGT |
| | R | CACAACTCCTCCAATCACAACT |
| OASL | F | GGGACAGAGATGGCACTGAT |
| | R | AAATGCTCCTGCCTCAGAAA |
| IFIT2 | F | ACGGTATGCTTGGAACGATTG |
| | R | AACCCAGAGTGTGGCTGATG |
| IFIT3 | F | CGGAACAGCAGAGACACAGA |
| | R | ATGGCATTTCAGCTGTGGA |
| DDX60 | F | AAGGTGTTCCTTGATGATCTCC |
| | R | TGACAATGGGAGTTGATATTCC |
| IFi6 | F | CTGTGCCCATCTATCAGCAG |
| | R | GGGCTCCGTCACTAGACCTT |
| SST1 | F | AACCACTGTGACGGGAGAAA |
| | R | CTGGGACAGGACGAGACAC |
| SATa | F | AAGGTCAATGGCAGAAAAGAA |
| | R | CAACGAAGGCCACAAGATGTC |
| HERVK | F | AGAGGAAGGAATGCCTCTTGCAG |
| | R | TTACAAAGCAGTATTGCTGCCCGC |

**Figure 1-figure supplement 1** 

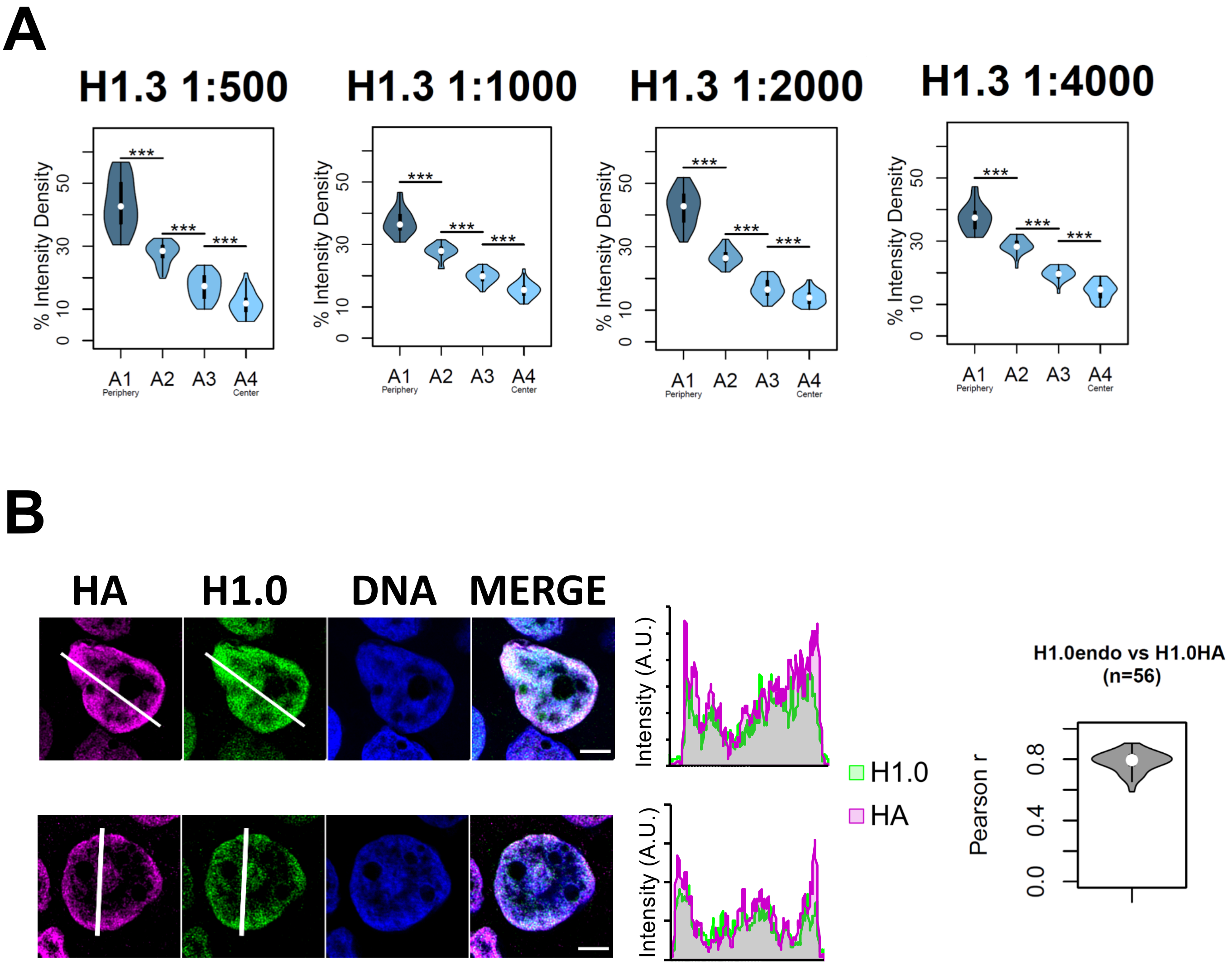

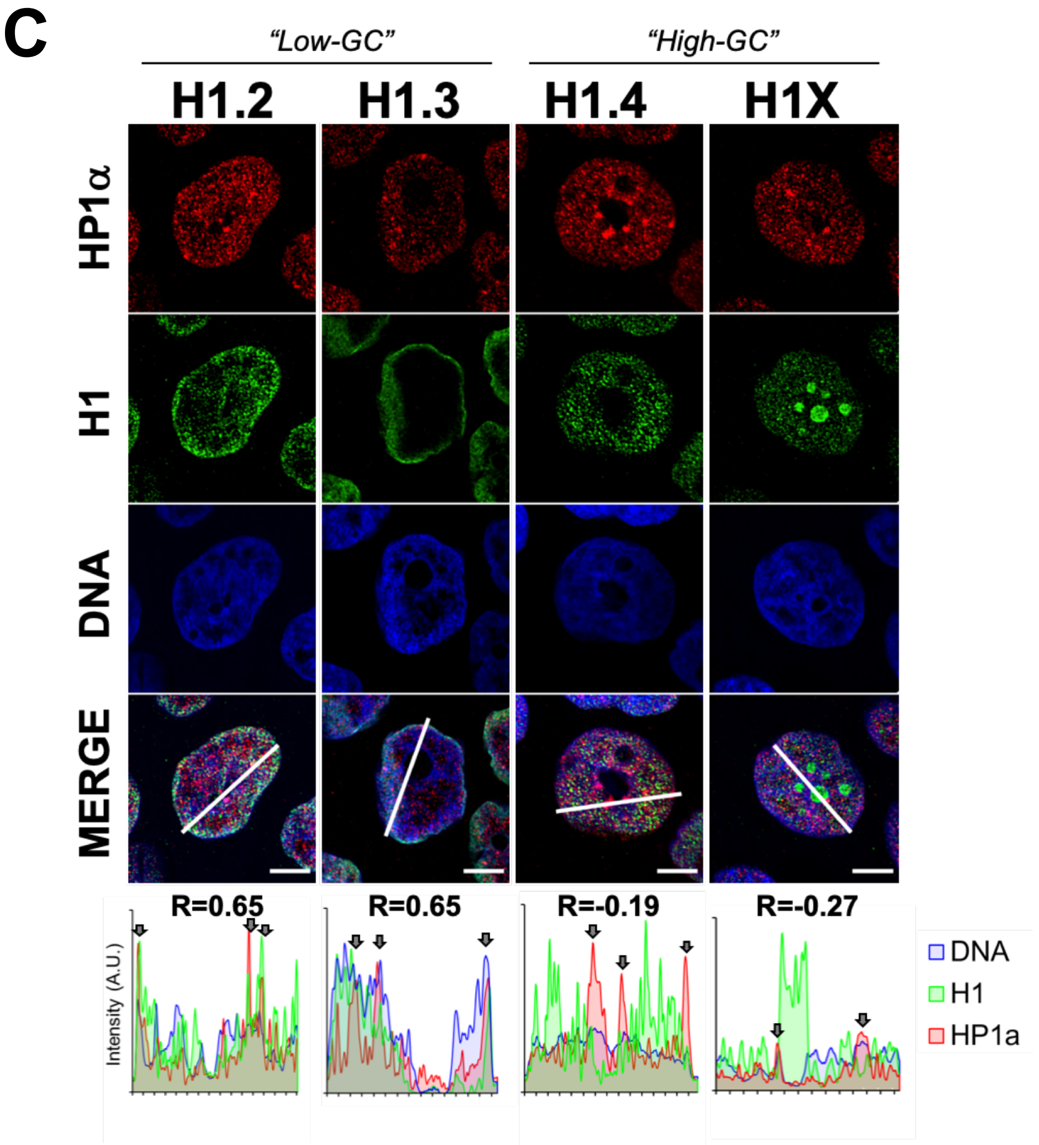
C
"Low-GC"
"High-GC"
H1.2
H1.3
H1.4
H1X
HP1α
H1
DNA
MERGE
R=0.65
R=0.65
R=-0.19
R=-0.27
Intensity (A.U.)
DNA
H1
HP1a

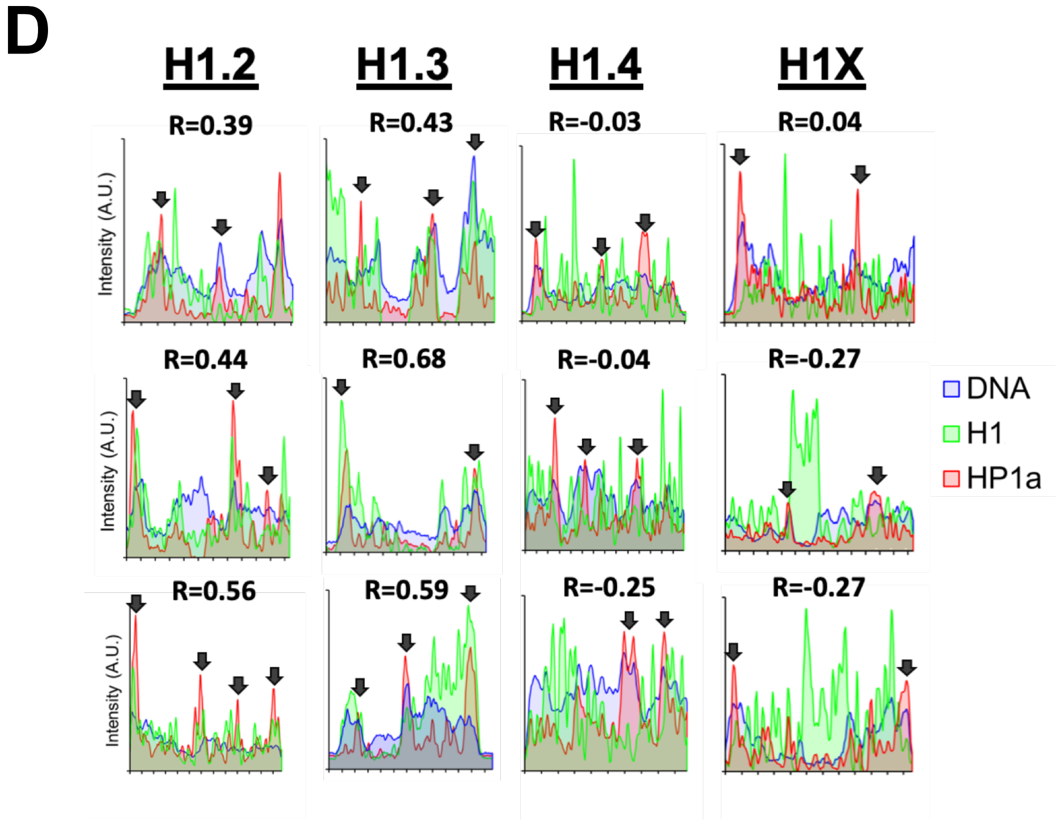
D
H1.2
H1.3
H1.4
H1X
R=0.39
R=0.43
R=-0.03
R=0.04
R=0.44
R=0.68
R=-0.04
R=-0.27
R=0.56
R=0.59
R=-0.25
R=-0.27
Intensity (A.U.)
DNA
H1
HP1a

**Figure 2-figure supplement 1** **Salinas et al.**

A

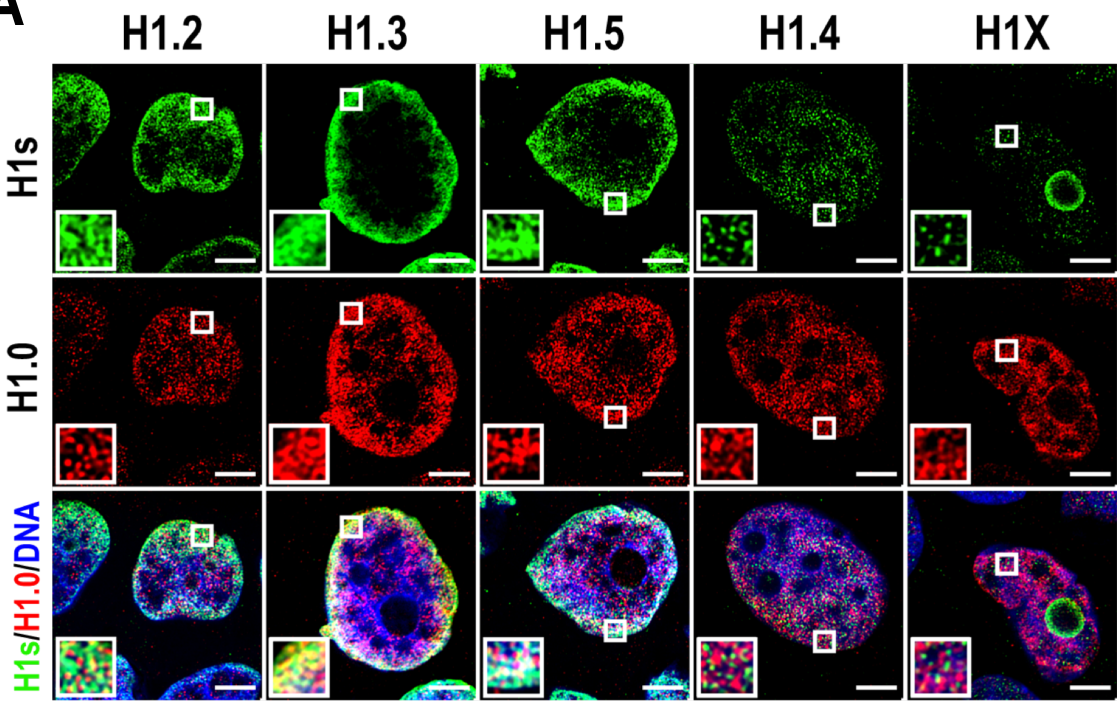


B

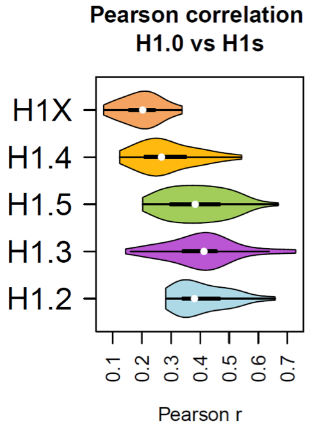


C

| ANOVA multiple comparison + Tukey multiple comparisons | |
|---|---|
| Histones | p-adj |
| H1.4-H1X | 0.0013 (**) |
| H1.5-H1X | 0 (***) |
| H1.3-H1X | 0 (***) |
| H1.2-H1X | 0 (***) |
| H1.5-H1.4 | 0.00002 (***) |
| H1.3-H1.4 | 0.000002 (***) |
| H1.2-H1.4 | 0.0000009 (***) |
| H1.3-H1.5 | 0.98 (ns) |
| H1.2-H1.5 | 0.95 (ns) |
| H1.2-H1.3 | 0.99 (ns) |

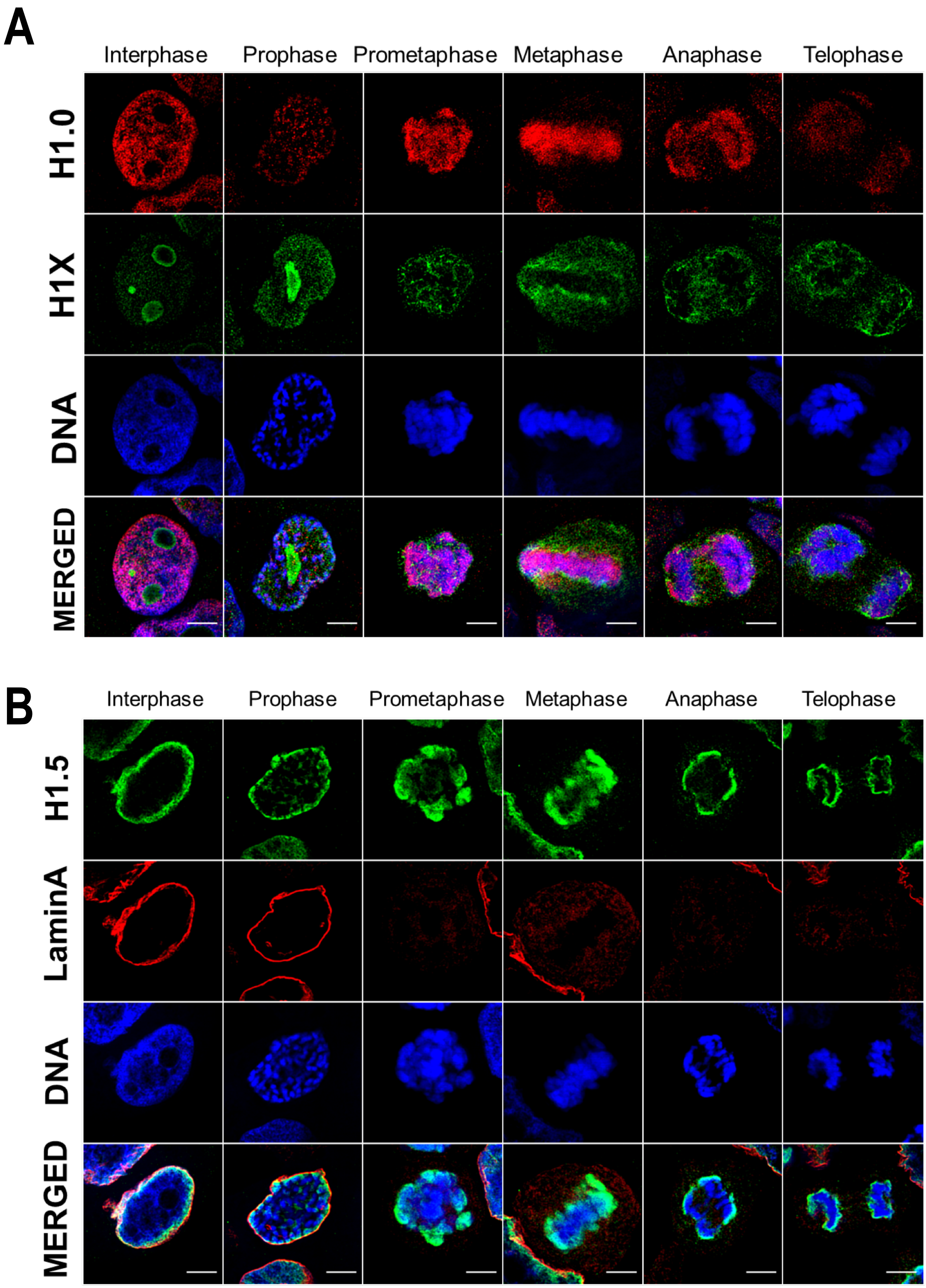
A
Interphase
Prophase
Prometaphase
Metaphase
Anaphase
Telophase
H1.0
H1X
DNA
MERGED
B
Interphase
Prophase
Prometaphase
Metaphase
Anaphase
Telophase
H1.5
LaminA
DNA
MERGED

C

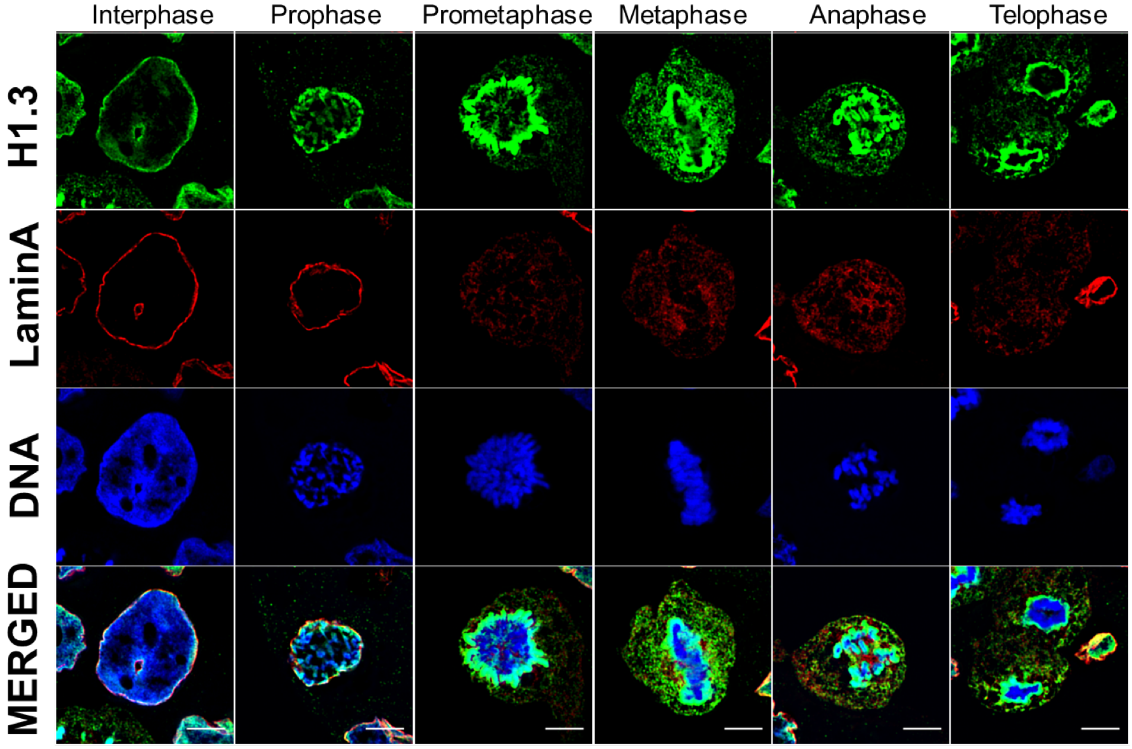
Interphase
Prophase
Prometaphase
Metaphase
Anaphase
Telophase
H1.3
LaminA
DNA
MERGED

D

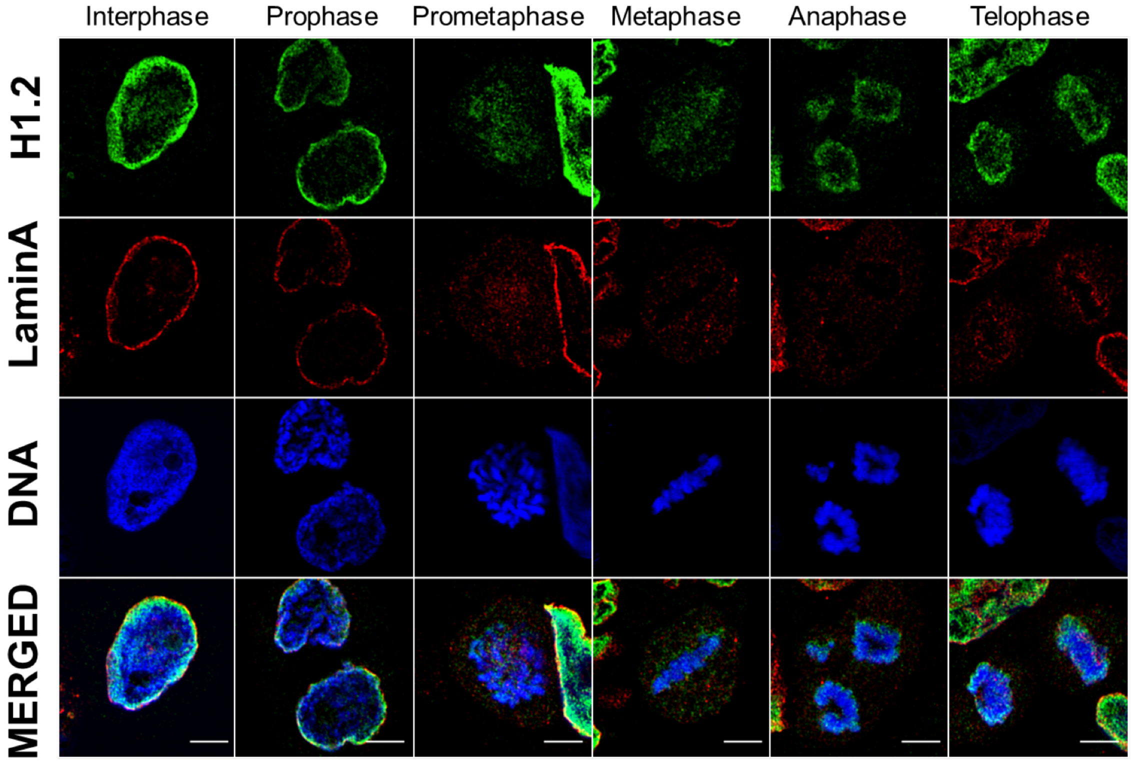
Interphase
Prophase
Prometaphase
Metaphase
Anaphase
Telophase
H1.2
LaminA
DNA
MERGED

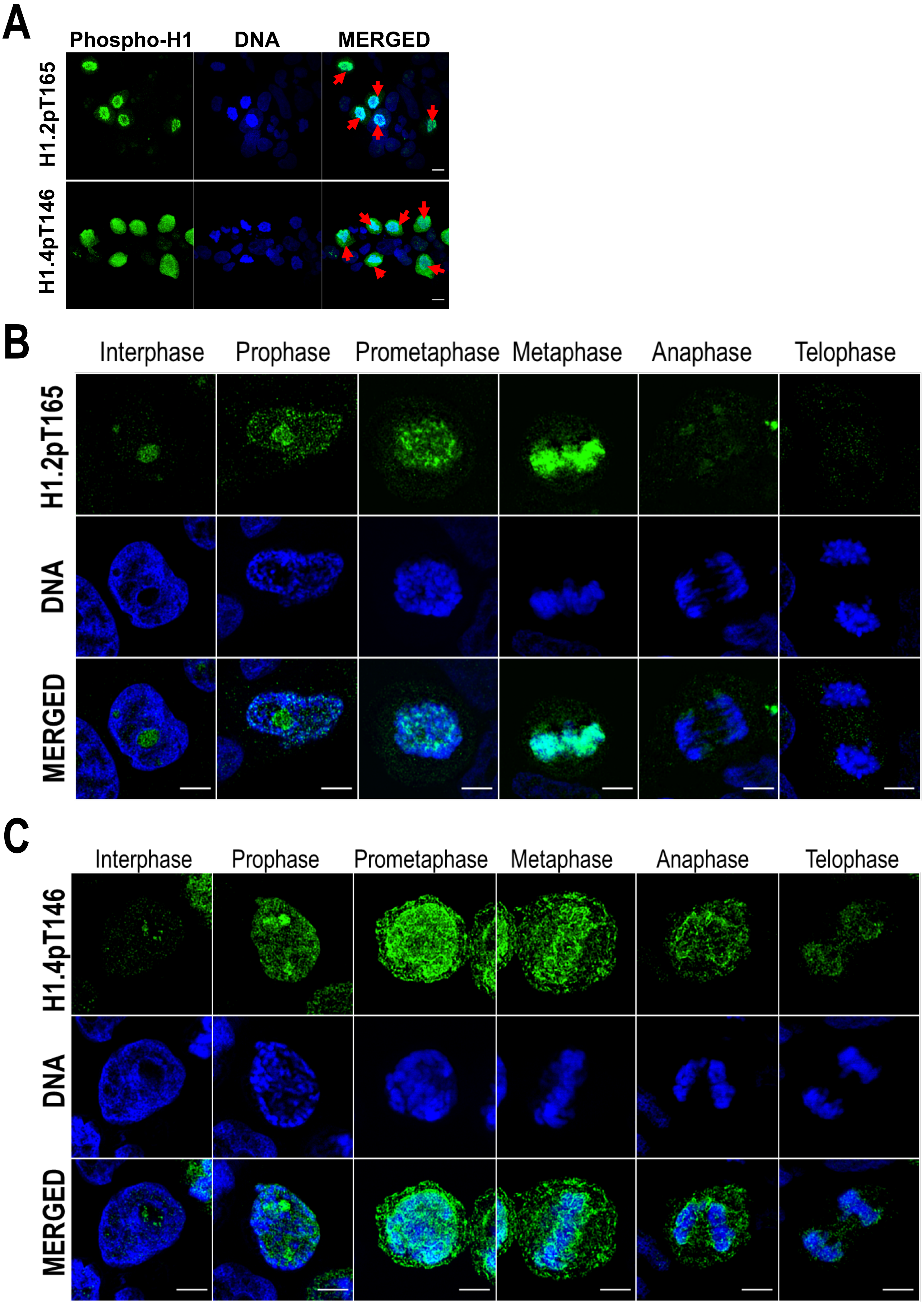
A
Phospho-H1
DNA
MERGED
H1.2pT165
H1.4pT146
B
Interphase
Prophase
Prometaphase
Metaphase
Anaphase
Telophase
H1.2pT165
DNA
MERGED
C
Interphase
Prophase
Prometaphase
Metaphase
Anaphase
Telophase
H1.4pT146
DNA
MERGED

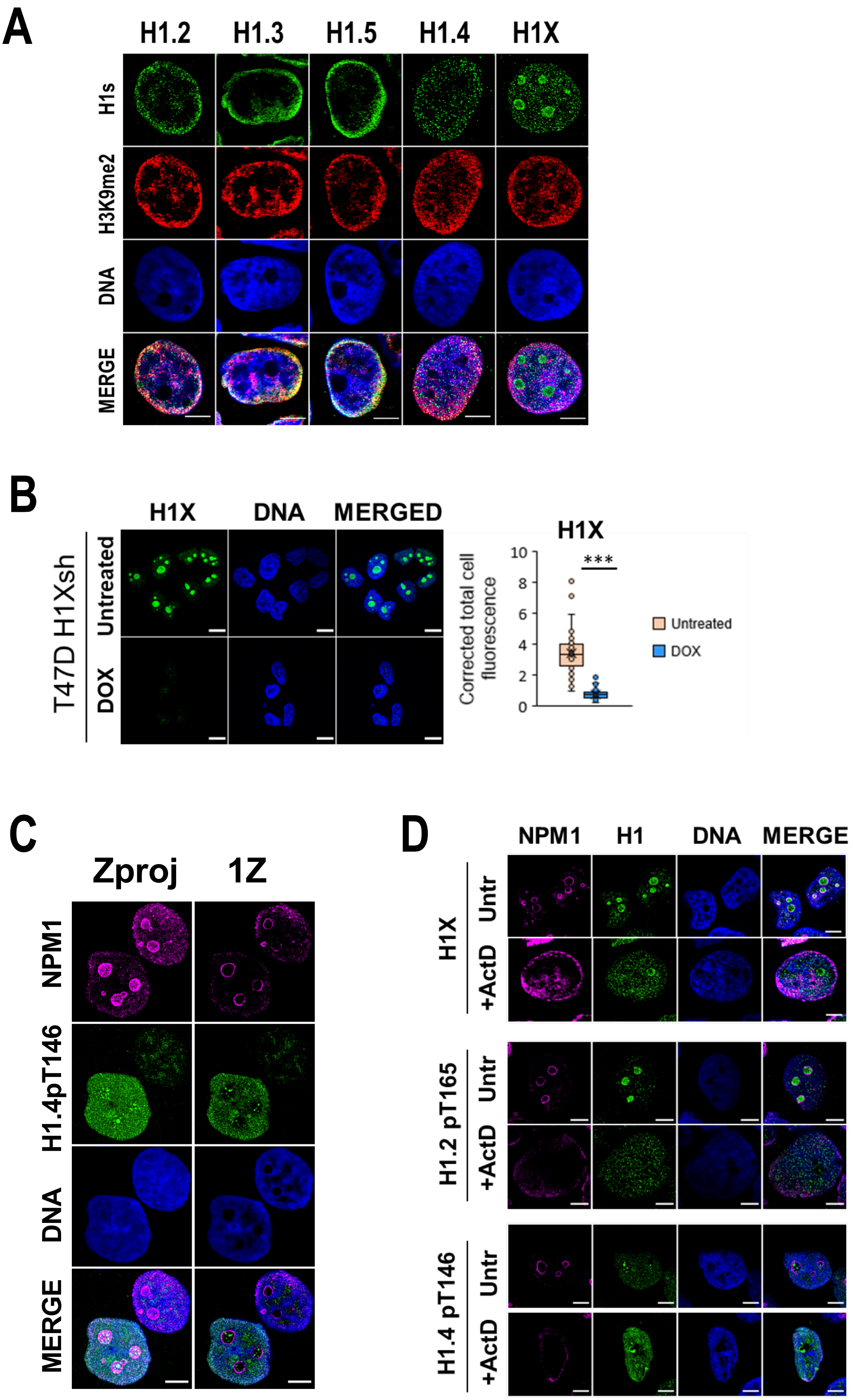
A
H1.2
H1.3
H1.5
H1.4
H1X
H1s
H3K9me2
DNA
MERGE
B
H1X
DNA
MERGED
T47D H1Xsh
Untreated
DOX
H1X
Corrected total cell fluorescence
10
8
6
4
2
0
***
Untreated
DOX
C
Zproj
1Z
NPM1
H1.4pT146
DNA
MERGE
D
NPM1
H1
DNA
MERGE
H1X
Untr
+ActD
H1.2 pT165
Untr
+ActD
H1.4 pT146
Untr
+ActD

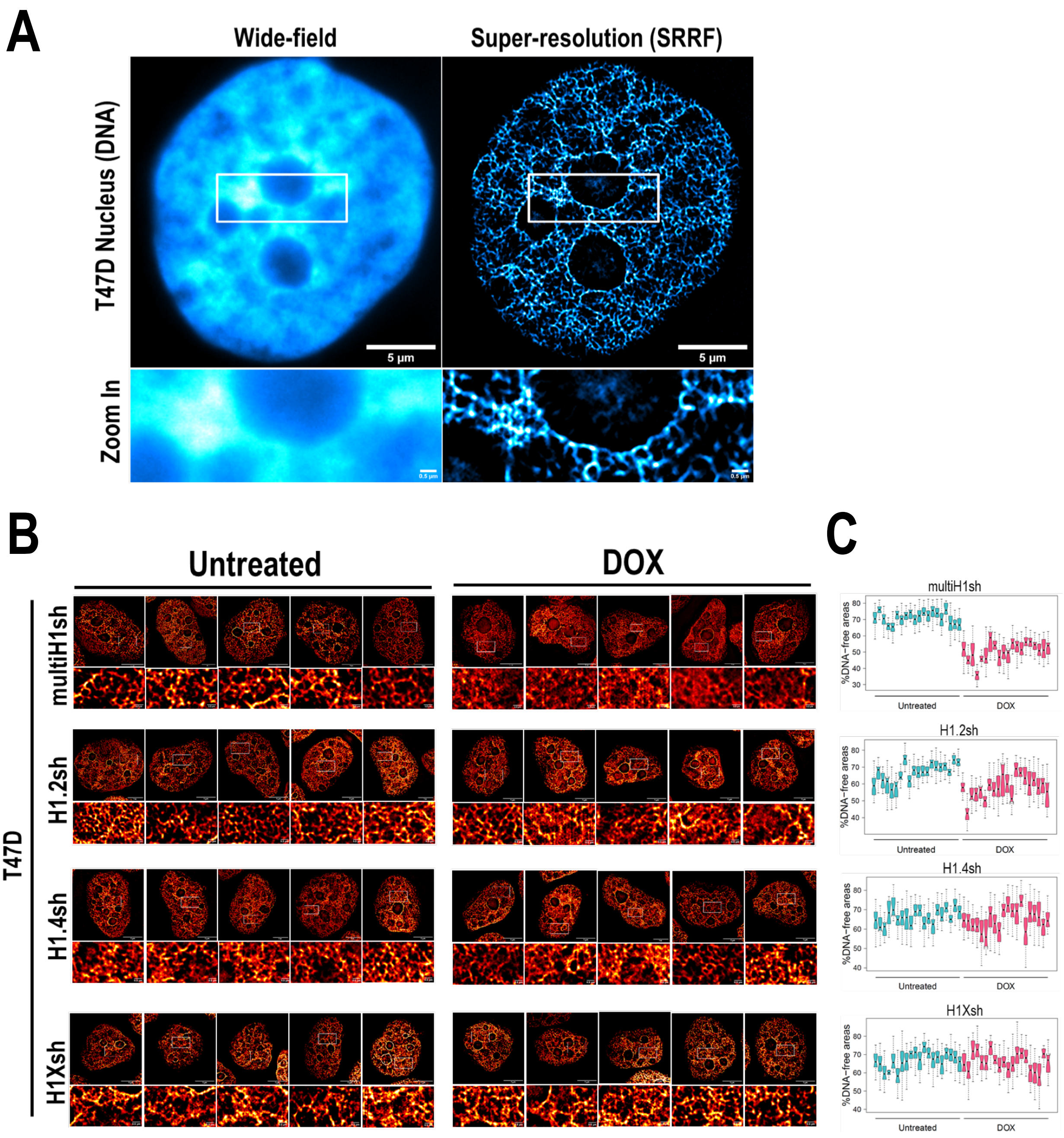
A
Wide-field
Super-resolution (SRRF)
T47D Nucleus (DNA)
5 µm
5 µm
Zoom In
B
Untreated
DOX
T47D
multiH1sh
H1.2sh
H1.4sh
H1Xsh
C
multiH1sh
H1.2sh
H1.4sh
H1Xsh
%DNA−free areas
Untreated
DOX

### Figure 5-figure supplement 1

Salinas et al.

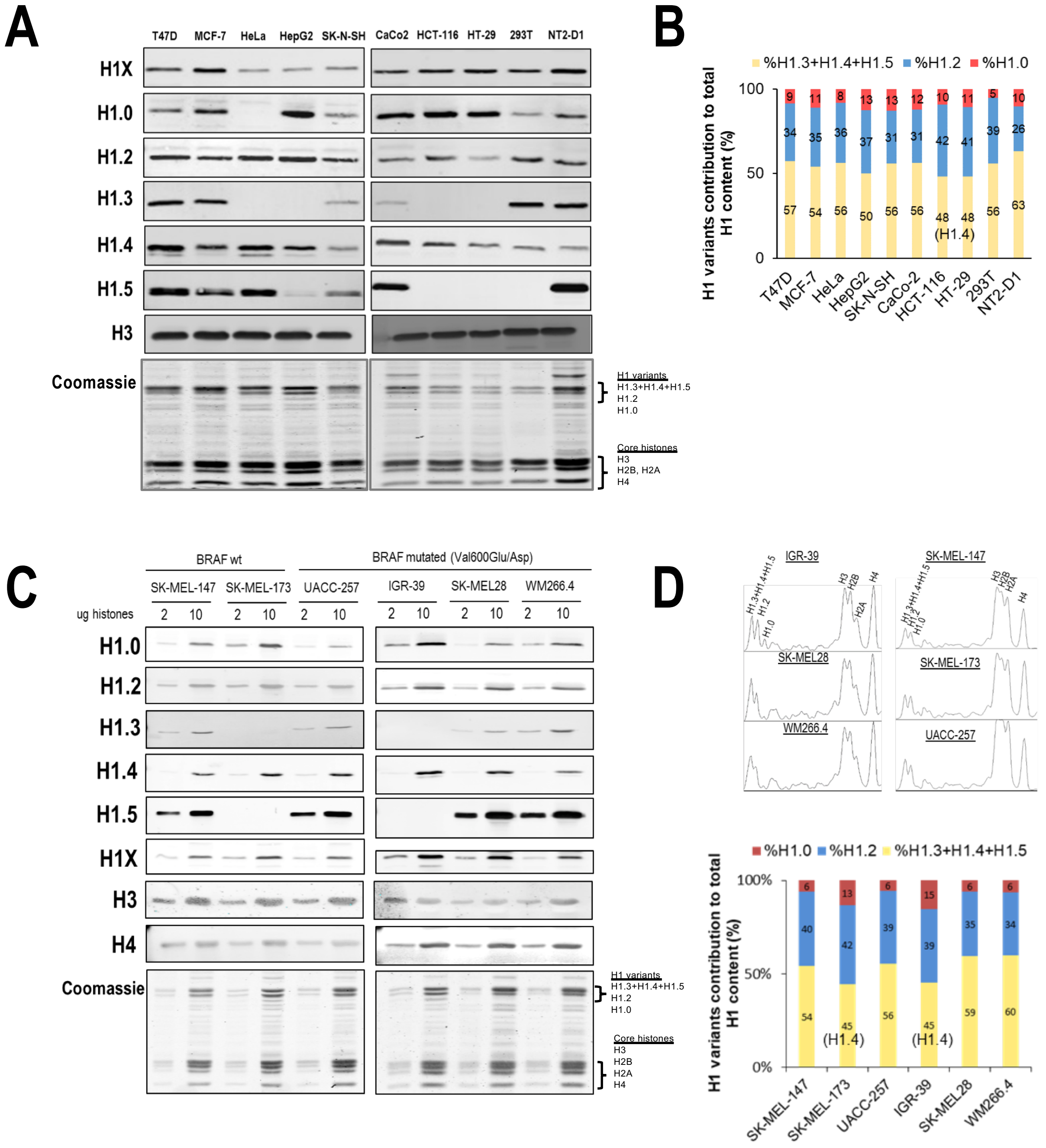


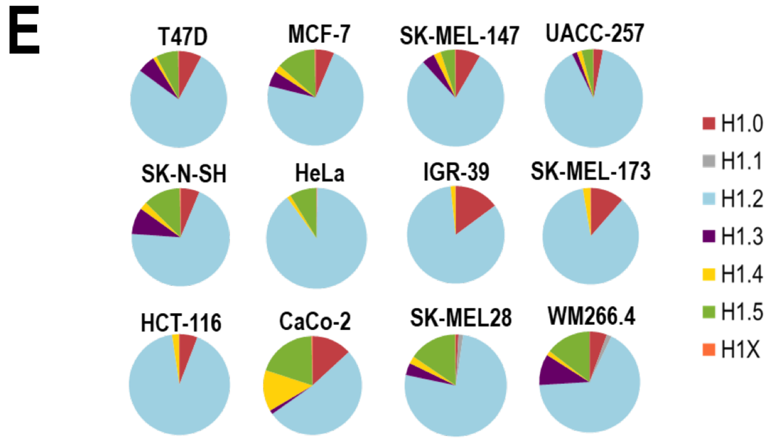


| (%) | H1.0 | H1.1 | H1.2 | H1.3 | H1.4 | H1.5 | H1X |
|---|---|---|---|---|---|---|---|
| T47D | 7.73 | 0 | 77.38 | 6.02 | 1.21 | 7.43 | 0.22 |
| MCF-7 | 6.28 | 0.01 | 72.59 | 5.22 | 2.38 | 13.25 | 0.28 |
| SK-N-SH | 6.26 | 0.01 | 69.83 | 8.82 | 2.54 | 12.27 | 0.28 |
| HeLa | 0.07 | 0.26 | 89.7 | 0 | 1.22 | 8.62 | 0.13 |
| HCT-116 | 5.82 | 0.01 | 91.81 | 0 | 2.24 | 0 | 0.12 |
| CaCo-2 | 13.14 | 0.02 | 52.07 | 1.29 | 13.53 | 19.6 | 0.37 |
| SK-MEL-147 | 8.36 | 0 | 79.91 | 4.26 | 2.45 | 4.83 | 0.19 |
| UACC-257 | 3.07 | 0 | 89.71 | 1.66 | 1.61 | 3.82 | 0.12 |
| IGR-39 | 14.82 | 0.07 | 83.35 | 0 | 1.65 | 0.01 | 0.1 |
| SK-MEL-173 | 11.39 | 0 | 85.97 | 0 | 2.51 | 0 | 0.13 |
| SK-MEL-28 | 0.83 | 1.53 | 75.94 | 3.87 | 2.37 | 15.32 | 0.14 |
| WM266-4 | 5.56 | 1.54 | 66.98 | 9.94 | 1.46 | 14.47 | 0.06 |

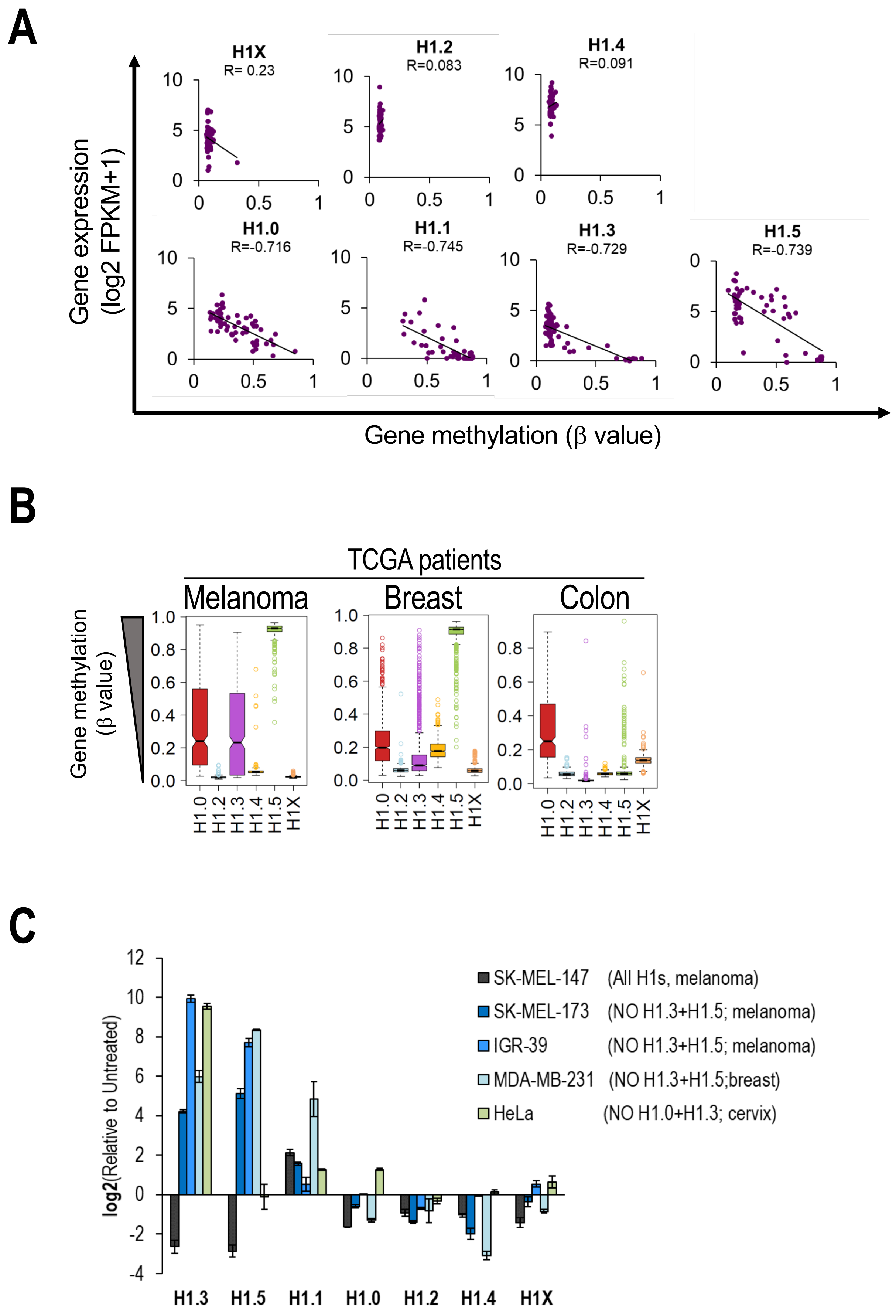
A
H1X
R= 0.23
H1.2
R=0.083
H1.4
R=0.091
H1.0
R=-0.716
H1.1
R=-0.745
H1.3
R=-0.729
H1.5
R=-0.739
Gene expression
(log2 FPKM+1)
Gene methylation (β value)
B
TCGA patients
Melanoma
Breast
Colon
Gene methylation
(β value)
H1.0 H1.2 H1.3 H1.4 H1.5 H1X
C
log2(Relative to Untreated)
SK-MEL-147 (All H1s, melanoma)
SK-MEL-173 (NO H1.3+H1.5; melanoma)
IGR-39 (NO H1.3+H1.5; melanoma)
MDA-MB-231 (NO H1.3+H1.5;breast)
HeLa (NO H1.0+H1.3; cervix)
H1.3 H1.5 H1.1 H1.0 H1.2 H1.4 H1X

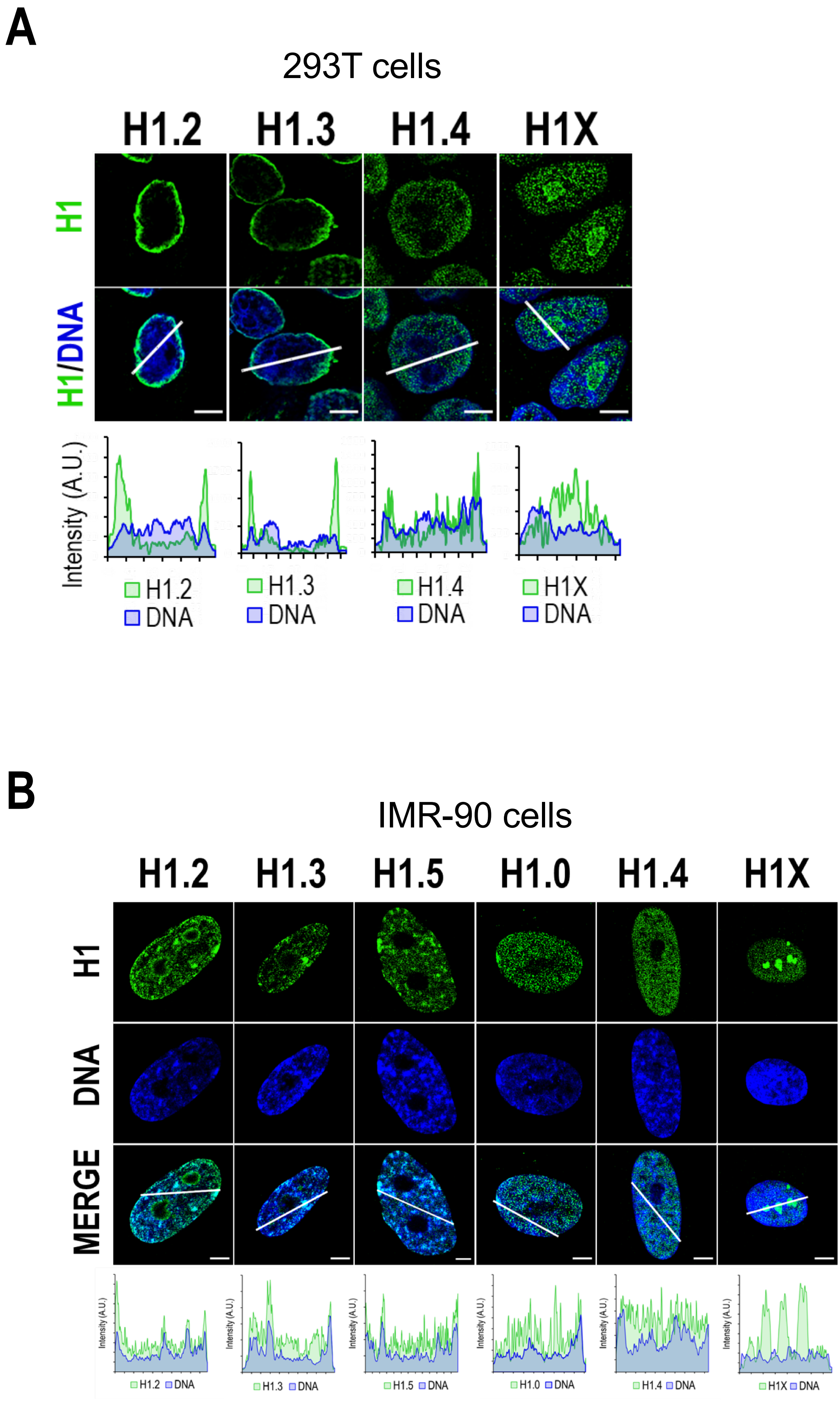
A
293T cells
H1.2
H1.3
H1.4
H1X
H1
H1/DNA
Intensity (A.U.)
H1.2
DNA
H1.3
DNA
H1.4
DNA
H1X
DNA
B
IMR-90 cells
H1.2
H1.3
H1.5
H1.0
H1.4
H1X
H1
DNA
MERGE
H1.2 DNA
H1.3 DNA
H1.5 DNA
H1.0 DNA
H1.4 DNA
H1X DNA

A

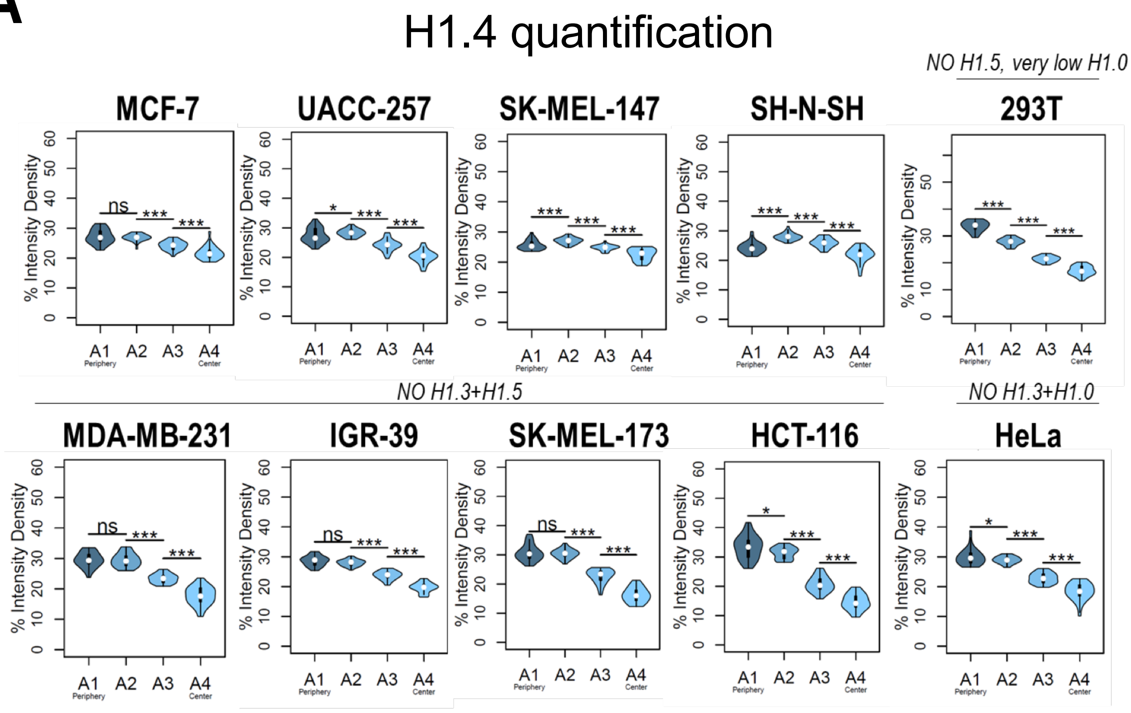
H1.4 quantification
NO H1.5, very low H1.0
MCF-7
UACC-257
SK-MEL-147
SH-N-SH
293T
% Intensity Density
ns
*
***
A1 Periphery
A2
A3
A4 Center
NO H1.3+H1.5
NO H1.3+H1.0
MDA-MB-231
IGR-39
SK-MEL-173
HCT-116
HeLa

B

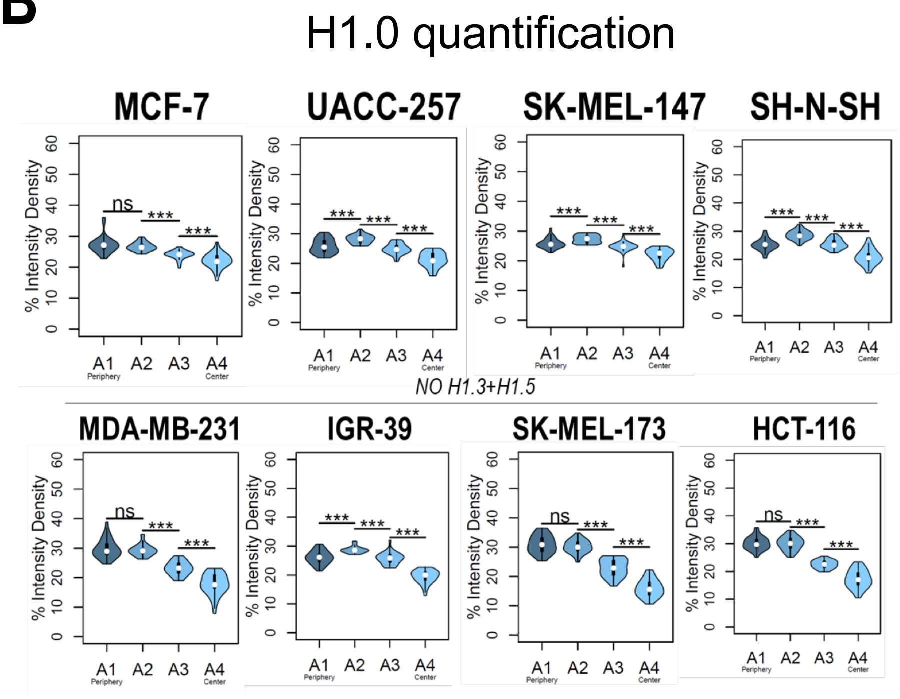
H1.0 quantification
MCF-7
UACC-257
SK-MEL-147
SH-N-SH
% Intensity Density
ns
***
A1 Periphery
A2
A3
A4 Center
NO H1.3+H1.5
MDA-MB-231
IGR-39
SK-MEL-173
HCT-116

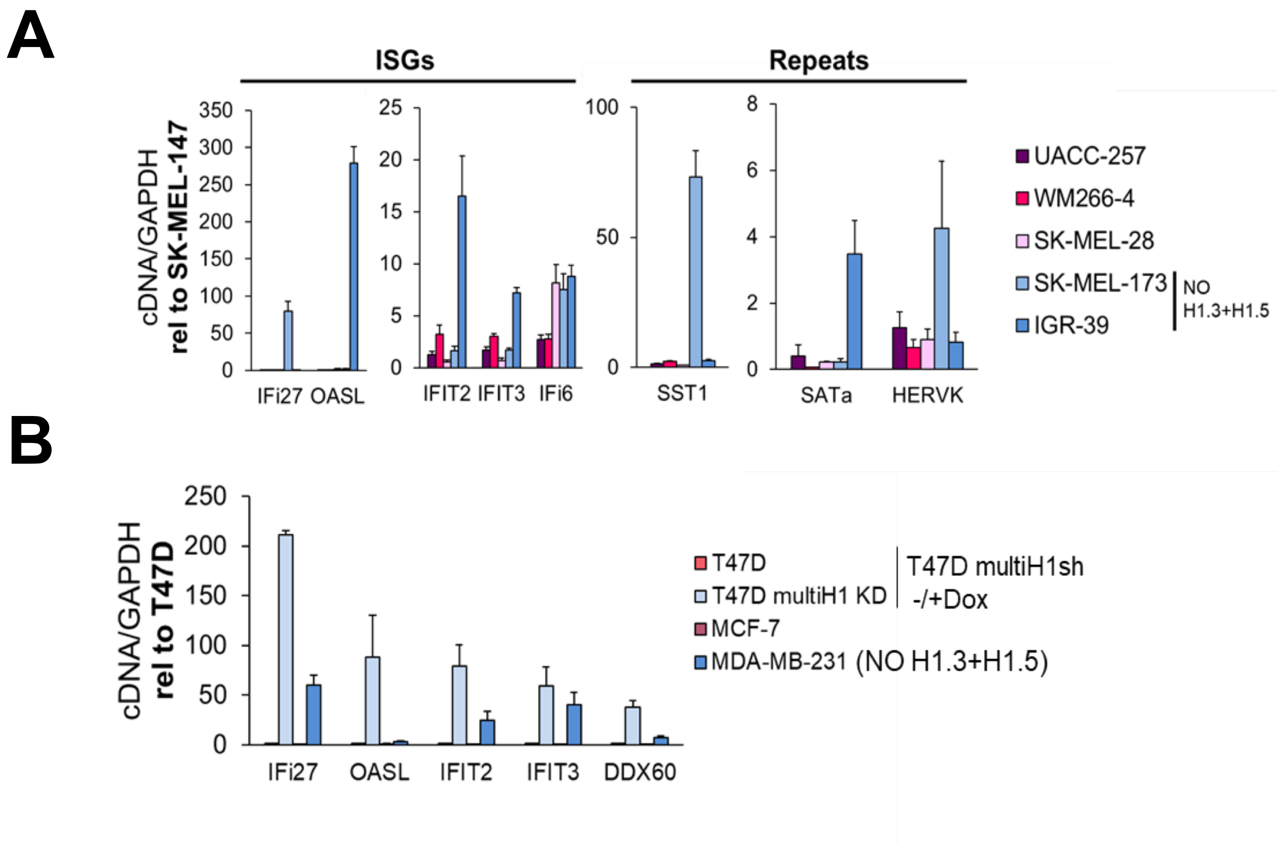
A
ISGs
Repeats
cDNA/GAPDH
rel to SK-MEL-147
350
300
250
200
150
100
50
0
IFi27 OASL
25
20
15
10
5
0
IFIT2 IFIT3 IFi6
100
50
0
SST1
8
6
4
2
0
SATa HERVK
UACC-257
WM266-4
SK-MEL-28
SK-MEL-173
IGR-39
NO H1.3+H1.5
B
cDNA/GAPDH
rel to T47D
250
200
150
100
50
0
IFi27 OASL IFIT2 IFIT3 DDX60
T47D
T47D multiH1 KD
T47D multiH1sh -/+Dox
MCF-7
MDA-MB-231 (NO H1.3+H1.5)
